# *i*BRAB: *in silico* based-designed Broad-spectrum Fab against H1N1 Influenza A Virus

**DOI:** 10.1101/2020.09.01.277335

**Authors:** Phuc-Chau Do, Trung H. Nguyen, Uyen T.M. Vo, Ly Le

## Abstract

Influenza virus A is a significant agent involved in the outbreak of worldwide epidemics, causing millions of fatalities around the world by respiratory diseases and seasonal illness. Many projects had been conducting to investigate recovered infected patients for therapeutic vaccines that have broad-spectrum activity. With the aid of the computational approach in biology, the designation for a vaccine model is more accessible. We developed an *in silico* protocol called *i*BRAB to design a broad-reactive Fab on a wide range of influenza A virus. The Fab model was constructed based on sequences and structure of available broad-spectrum Abs or Fabs against a wide range of H1N1 influenza A virus. As a result, the proposed Fab model followed *i*BRAB has good binding affinity over 27 selected HA of different strains of H1 influenza A virus, including wild-type and mutated ones. The examination also took by computational tools to fasten the procedure. This protocol could be applied for a fast designed therapeutic vaccine against different types of threats.

## Introduction

Influenza A virus is known as an agent causing seasonal illness of respiratory diseases. This agent is also responsible for the outbreak of worldwide epidemics, causing millions of fatalities worldwide. The first well-documented pandemic appeared to start in Russia and reached St. Petersburg in October 1889.[1] However, the H1N1 Spanish flu in 1918 is the nightmare pandemic, starting with 550,000 deaths and increasing to at least 50 million, possibly up to 100 million, of deaths worldwide.[2] Form then, many pandemics occurred caused by different types of influenza A virus, such as H7N9, H3N2 or H5N1, were recorded.[3–6] Despite different types of threats, the H1N1 from the 1918 pandemic is still the potential threat nowadays by its highly frequent shift of genetic components.[7] To fight back the influenza virus, doctors and scientists around the world are working on vaccines and drugs specifically designed for this type of disease. Most of the prophylactic vaccine developed in the past and currently used focus on neutralizing Hemagglutinins on the surface of a specific strain of influenza A virus.[8] In parallel with research to improve the prophylactic vaccine’s effectiveness against influenza A viruses, many research groups worldwide work on antibodies collected from recovered infected people for therapeutic vaccines. The therapeutic vaccine encourages the infected body to fight against the disease harder.[9] Antibodies are widely used as therapeutic vaccines due to their high specificity and low toxicity. Also, they can neutralize the viral and signal the macrophages for viral degradation. Broad-reactive antibodies against many types of influenza A virus are studied extensively to fight against new emergent incidents. The method applied in these studies is screening the available antibodies collected from recovered infect-patients from previous pandemics. However, the vast number and diversity of antibodies in one human body makes screening challenging to achieve and time-consuming. There are four big groups of researchers working in this field and giving some antibody (Ab) or antigen-binding fragment (Fab) candidates. Research groups initiated from James E. Crowe Jr. (Vanderbilt University School of Medicine) and Ian A. Wilson (The Scripps Research Institute) at the United States of America made the most reports about this therapeutic vaccine.[10–30] Another group from medical schools in the U.S.A led by Stephen C. Harrison, Barton F. Haynes, and Aaron G. Schmidt also provided a different list of candidates.[31–39] European and Chinese researchers also contributed to the working on the influenza-therapeutic vaccine. While Steven J. Gamblin, John J. Skehel and Antonio Lanzavecchia have been leading a group in Europe,[40–43] another Chinese group led by Linqi Zhang and Xinquan Wang, who had studied several candidates.[44–49]

Antibodies, or immunoglobulins, are heterodimeric proteins comprised of two heavy chains and two light chains that eventually create a “Y” shaped molecule. The activity of Ab is determined by the tip regions, which are called variable regions giving each Ab its specific binding activity against the antigen (Ag).[50, 51]. The variable regions are further divided into hypervariable and framework regions. Within a Fab formed by a complete light chain and a portion of a heavy chain, there are six hypervariable regions (three for each chain), and they have a high ratio of different amino acids within regions. The hypervariable regions, in some cases, are preferred as Complementarity-determining regions (CDRs).[52] During the binding of Fab to HA, these CDRs act as a “key” to “lock” the receptor-binding site. Although CDRs of an individual Ab binds a limited and defined set of ligands, it still can bind to a set of practical ligands sharing little or no similarity.[53] Besides, a single variable fragment of Fab can be genetically engineered to represent the specific property of parent Ab against antigens.[54]

In recent years, bioinformatics technology has been used globally to address disease issues and become a powerful tool for rational drug design. By studying a wide range of broad-spectrum Fabs and helping with the computational approach, we developed a protocol called *i*BRAB (***i**n silico* **B**road-**R**eactive **A**ntigen-**B**inding fragment) to design potential therapeutic vaccines against a wide range of influenza A virus. In this protocol, Abs/Fabs having broad-spectrum activity against a wide range of trains are computationally investigated their sequences and 3-dimension structures to propose a Fab model. This proposed model can have broad-reactive on different strains of influenza A virus and overcomes current products’ limitation over mutations of antigens.

## Models and Methods

### Models

Sequences and 3D structures of broad-spectrum antibodies (Ab) or antigen-binding fragment (Fab) were used as templates in this research. The information was collected from the Research Collaboratory for Structural Bioinformatics – Protein Data Bank (RCSB-PDB) database. Between 2010 and 2019, 63 published Abs and Fabs can bind and neutralize influenza hemagglutinin (HA). Ten of them have binding at the receptor-binding site (RBS) and neutralizing activities on wide-range strains of the same H1 subtype. Both sequence and 3D structure of the heavy or light chain of each Ab/Fab were extracted into separated files for later analysis. Similarly, in the RCBS-PDB database, among 93 available HA proteins belonged to the H1 subtype, 27 proteins were selected and refined to exam the binding affinity of the proposed Fab model on HA protein of different H1N1 strains.

### Multiple sequence alignment (MSA)

The alignment of multiple heavy/light-chain sequences was done by the MEGA-X program to propose a sequence of broad-spectrum Fab model.[78] Heavy chains and light chains were aligned separately using MUltiple Sequence Comparison by Log-Expectation (MUSCLE). The parameters were set with Gap open −2.9, Gap extends 0.0, and Hydrophobicity multiplier 1.2.

### Complementarity-determining region (CDR) determination

CDRs of chains were examined on IMGT/3Dstructure-DB and IMGT/2Dstructure-DB database.[79–81] For several newly published Abs/Fabs, the data have not been available on the IMGT database yet, Paratome tool was used to identify the CDRs.[52] The positions of CDRs in heavy and light chains were recorded and marked on the multiple sequence alignment results.

### Superimposition

All heavy chains and light chains of Abs/Fabs were run with UCSF Chimera 1.13.1 for the superimposition approach.[82] Smith-Waterman alignment algorithms, BLOSUM-62 matrix with Gap extension penalty 1 were used for alignment. The secondary structure was included and accounted for 30% of the total score. The matching process was iterated by pruning long atom pairs until no pair exceeds 2.0Å. Then, all chains were chosen for alignment with residue-residue distance cut-off 5.0Å; the aligned residues were put in a column if they are within the cut-off value of at least one other structure. Circular permutation and iterate superimposition/alignment were turned on with 3 times iterated alignment and 3 consecutive columns in stretches of superimposing full columns.

### 3D-structure modeling

Based on the superimposition from UCSF Chimera, sequences of heavy and light chains for the Fab model were proposed for the Fab model against the H1 subtype. 3D structures of proposed models were constructed by MODELLER version 9.21.[83] All ten Abs/Fabs from previous experiments were used as templates for the modeling. Some parameters were kept as a previous superimposed experiment, while the gap extension penalty was increased to 3 to reduce the gap introduction. A hundred new 3D models for each query were built and evaluated by Discrete optimized protein energy (DOPE) score.[84] The output model, which generated a minimum molecular probability density function (molpdf) value and the DOPE score, was continued with ab-initio loop refinement at HCDR3 and LCDR3 regions, and regions with DOPE peak higher than −0.02. The refinement was done for a hundred outputs.

### Flexible docking

Molecular docking is an effective method for virtual screening.^87^ High Ambiguity Driven protein-protein DOCKing (HADDOCK) 2.4 server was used as the flexible docking simulation between the Fab model and HA proteins to estimate the binding affinity.[85, 86] All amino acids at CDR regions of the Fab model and RBS region of HA were chosen as active residues without penalty. All other parameters were set at default. RMSD-based and contact-based clustering was done with cut-off values 7.5Å and 0.75, respectively. The algorithm of Daura et al. and Rodrigues et al. was used separately in each clustering. The binding pose with the lowest binding energy of each model-HA complex was extracted to preliminary evaluate the antibody efficiency on different receptors.

### Protein-protein interaction (PPI) calculation

The PPI analysis was performed by BIOVIA Discovery Studio 2020 (Dassault Systèmes, San Diego, California, USA). A web-based server Protein Interaction Calculator of the Molecular Biophysics Unit, Indian Institute of Science (Bangalore), was used together to maximize the analysis.[87] The complex systems were examined all types of hydrogen bonds, hydrophobic interaction, electrostatic interaction, and steric bumps intermolecularly between the Fab model and HA proteins.

### Statistical analysis

All the graphs, together with mean, standard deviation, and other parameters shown in figures, tables were drawn and calculated by package tidyverse in R tool (with interface RStudio software). P-value was calculated using the one-way ANOVA method – Tukey pairwise test, and statistical significance was set at p-value < 0.05.

## Results and Discussion

### The constant regions of heavy chains are the same through ten Fab templates, while the constant regions of light chains are in both λ and κ groups

To examine the Fab templates in amino acid sequence and 3D structure, we did MSA by the MUSCLE algorithm of the MEGA-X program and superimposition using the UCSF Chimera 1.13.1 tool. The results show that amino acid sequences at the framework and constant regions have higher conservative property, while variable regions vary between Fab templates (Fig 1). In the constant regions of Fab templates, both MSA and superimposition give consistent results with each other. The sequences and 3D structures are preserved through ten heavy chains and two groups of light chains. There is only one difference at 130-loop forming between beta-sheets, although the heavy chains’ amino acid sequences are the same. This incident is due to the biophysical activity in the physiological environment that loop structure can be flexible cross-linking with surrounding chemical molecules.[88, 89] In additional, despite the difference in amino acid sequences of λ and κ light chains, the scaffold of 3D structures is similar with beta-sheets and alpha-helices, and the formed loops in κ are longer than in λ light chains. Many works have shown that as loop length increases, the loop’s stability is decreased by thermal and chemical denature.[90, 91] With shorter loops, the structure of a protein is more stable in the physiological state; thus, the λ light chain is preferable in our proposed broad-spectrum Fab model. Besides, this λ light chain can help in avoiding antigen-specific selection by the pre-B cell.[92] MSA results show a very divergence between Fab templates in both heavy and light chains in the variable regions. However, the 3D structure of framework segments at these variable regions are superimposed well. These segments are essential to form the scaffold for Ab and generate three CDR loops. As expected, 3D structures of CDR loops between frameworks are profoundly different between Fab templates. The length of loops, presence of helix-turn in CDR1 and CDR3, or the extension of beta-sheet near CDR3 could make the difference. The HCDR1 loop forms small alpha-helix in 2D1, 5J8, 1F1, CH67, and H2897 Fabs, while others have big-angle HCDR1 loops. Moreover, HCDR2 loops, which are relatively short and made by two beta-sheets near the HCDR1, give loops a semi-flexible structure. These criteria explained the minor interaction of HCDR1 and HCDR2 in Fab binding on the surface of HA. Notably, the HCDR3 loop, which is the primary interaction with HA and reported as a key to insert into the RBS to prevent the virus from accessing sialic acid on the host cell membrane, has transitioned with different influenza A strains differently. While CH67, H1244 and H2227 Fab templates have alpha-helix turn in HCDR3, 5J8, 6639, and 641 I-9 Fabs have longer beta-sheet, and others have a bigger loop with a long sequence. The three CDR loops in light chains have preserved 3D structure within their types. The λ light chains have alpha-helix at CDR2 and long CDR3 loop, while κ light chains have shorter CDR3 loop and no alpha-helix at CDR2, which can be explained by the MSA results between them.

**Fig 1.**
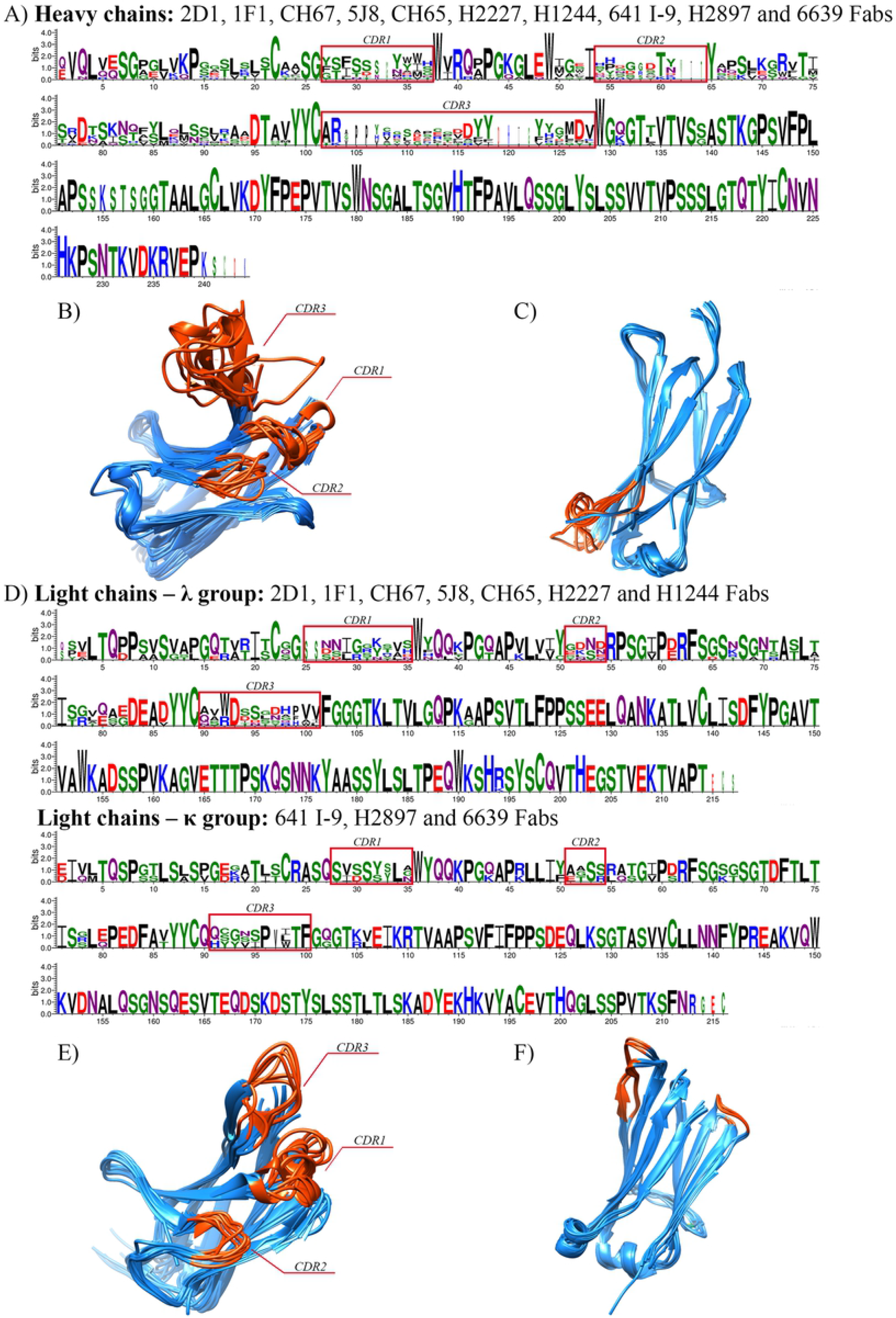
Multiple sequence alignment (MSA) and superimposition of ten heavy/light chains of broad-spectrum Fab templates against the H1 influenza A virus. MSA of (A) heavy chains and (D) light chains was done by the MEGA-X program. The amino acids’ logo was drawn using WebLogo version 3.7.4 with auto background composition and colored by chemical properties: green – polar, purple – neutral, blue – basic, red – acidic, and black – hydrophobic amino acids. 3D-structure superimposition of variable regions (B and E) and constant regions (C and F) of heavy and light chains, respectively, as illustrated by UCSF Chimera 1.13.1 and colored by the difference in 3D conservation of structure: blue – high and orange – less conservation.

### Proposed sequence of broad-spectrum Fab model against H1 influenza A virus includes different importance interacted amino acids of templates

Framework and constant regions have a minor role in binding between antibody and antigen,[53] thus, the previous MSA and superimposition results confirm the amino acid sequence of these regions for the Fab model. Essential amino acids of each Ab/Fab templates, which have crucial interaction with HAs, were considered when the model’s sequences were proposed. All the critical interactions were collected and distributed to the proposed sequence to increase the possible interactions with HA (Table 2). Many works had proven that mutations at RBS could abolish the binding between Fab and HA by deleting the interaction. One example, the mutation of Lys166 to Asp166 can cause the deletion of a salt bridge between Lys166 and Asp93L in 2D1 Fab, thus, the addition of other amino acids next to that position which can form interaction with Asp could increase the binding activity with mutated strains.[17] Moreover, the long HCDR3 loop with a narrow-angle is preferable to form a distinct pitch to penetrate the RBS region, reaching the deepest position.[32] In previous studies, there was no report on detailed interactions formed between HCDR1 and HCDR2, even they were mentioned as binding factors with HAs. Most of the HA interactions were reported and gathered in the HCDR3 region. These interactions are formed with polar and hydrophobic amino acids at HCDR3, which intruded into RBS. In detail, Arg and Asp are dominant in the collection, whereas the other polar and hydrophobic amino acids vary on different Fab templates. Many essential amino acids were combined in the model to increase the numbers of formed interactions. Aspartate amino acids were noted on the model’s heavy chain sequence at position 100, 101, 108, and 111. This amino acid interacts with different amino acids of HA. In 5J8, Asp100^H^ has an interaction with Trp153 and Leu194. Asp106^H^ of 641 I-9 makes interaction with Ala137 and Ser145. Asp107^H^ of CH65 and CH67 Fabs take part in Ser136, Ala137, Arg226 interactions. Furthermore, Asp110^H^ of H1244 and H2227 show a link with Ile194.[12, 32, 35, 36, 38] For positions 99 and 102, Arg100^H^ of H2897 and Arg103^H^ of 6639 were selected, which fixes HCDR3 conformation and makes extensive interaction with Tyr98, Thr136, Asp225, Arg226 in HA, respectively.[31, 55] For others, Val102^H^, Gly103^H^ and Thr105^H^ of H2897; Glu104^H^ of 6639 Fab; Val106^H^ of CH65 and CH67; Pro109^H^ of H1244 and H2227, these amino acids are given for 104-107 and 110.[31, 32, 36, 38, 93] In H1244 and H2227 Fabs, the HCDR3 loop conformation is strengthened by the disulfide bridge between Cys103^H^ - Cys108^H^, which lets the loop easily insert into the RBS.[32] These Cys are also put in the model at 103 and 109-positions. Although the LCDR loops are not the primary interaction with HA, they are essential in making complement binding with other regions around the binding site. It is reported that aspartic acid is the crucial interactor, which is mentioned and identified in protein complexes of HA-32D6, HA-2D1, and HA-5J8.[12, 17, 94] This amino acid was added to position 54 and 96. The interaction between 32D6 and HA also emphasized the interaction between Tyr31^L^, Ala32^L^, Thr94^L^, Trp95^L^, and Ala97^L^ with Gly172, Asn173, Gln206, and Gln210. These amino acids were recruited to the model to maximize the interaction.[94] Tyr32^L^ and Tyr33^L^ of H2897 make similar linkage as Tyr31^L^ of 32D6; thus, one Tyr was added for position 31.[31] Gly29^L^ in 5J8 was also noted to pair with Asp54L in the Fab model to strengthen the interaction with Lys145.[12]

**Table 1.**
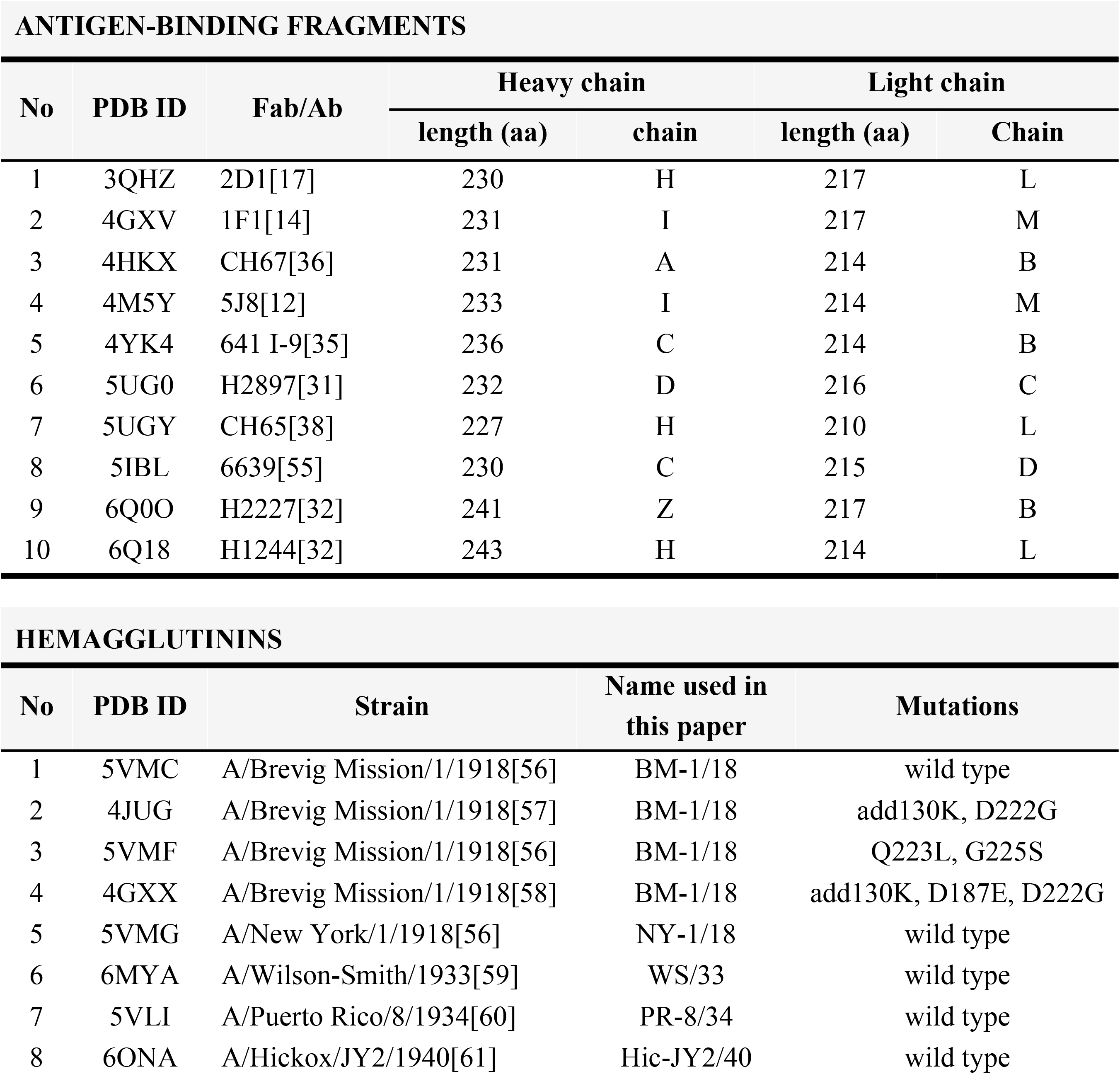
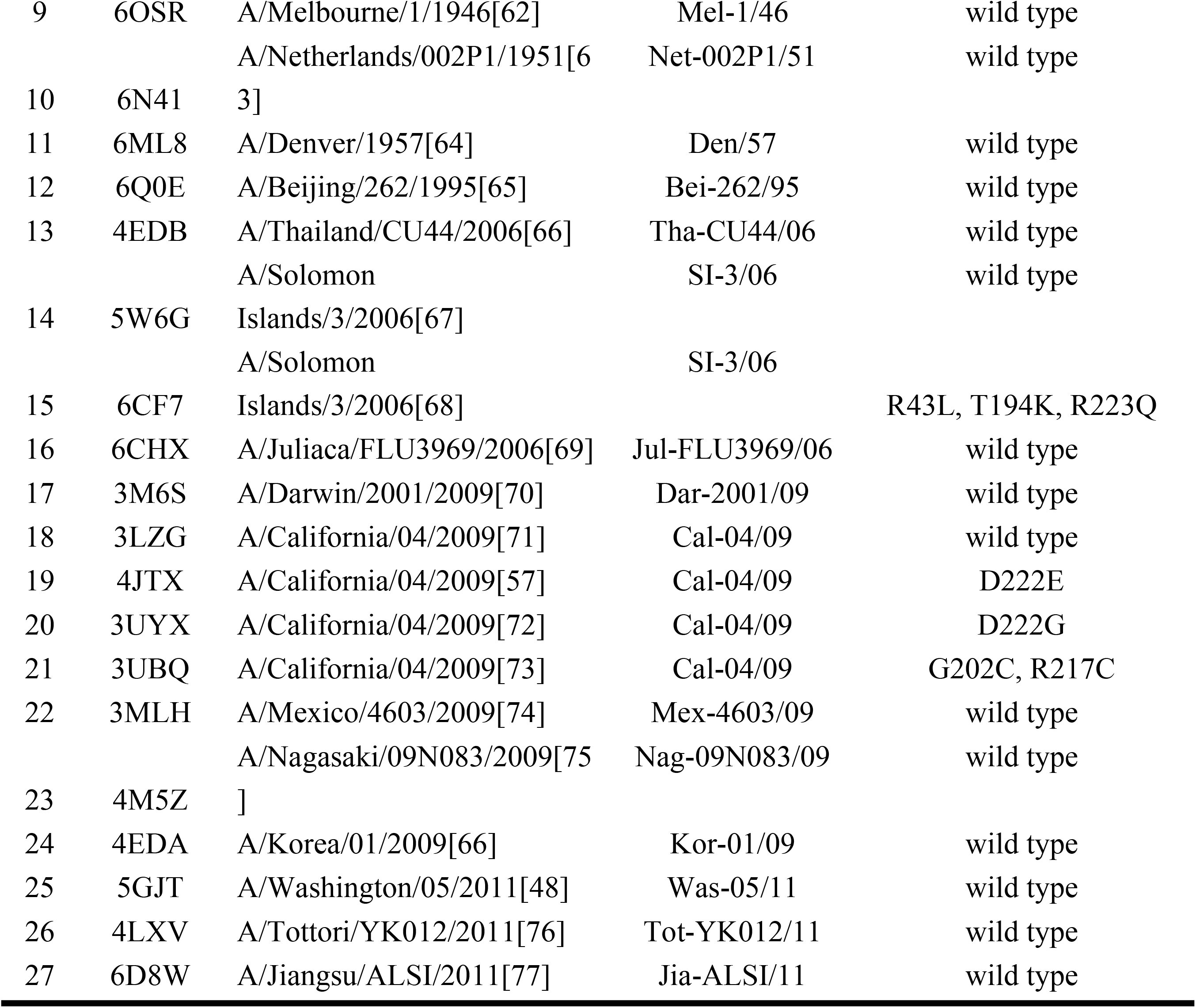
List of Broad-spectrum antibodies (Abs)/antigen-binding fragments (Fab) and Hemagglutinin (HA) proteins used in the experiment.

**Table 2.**
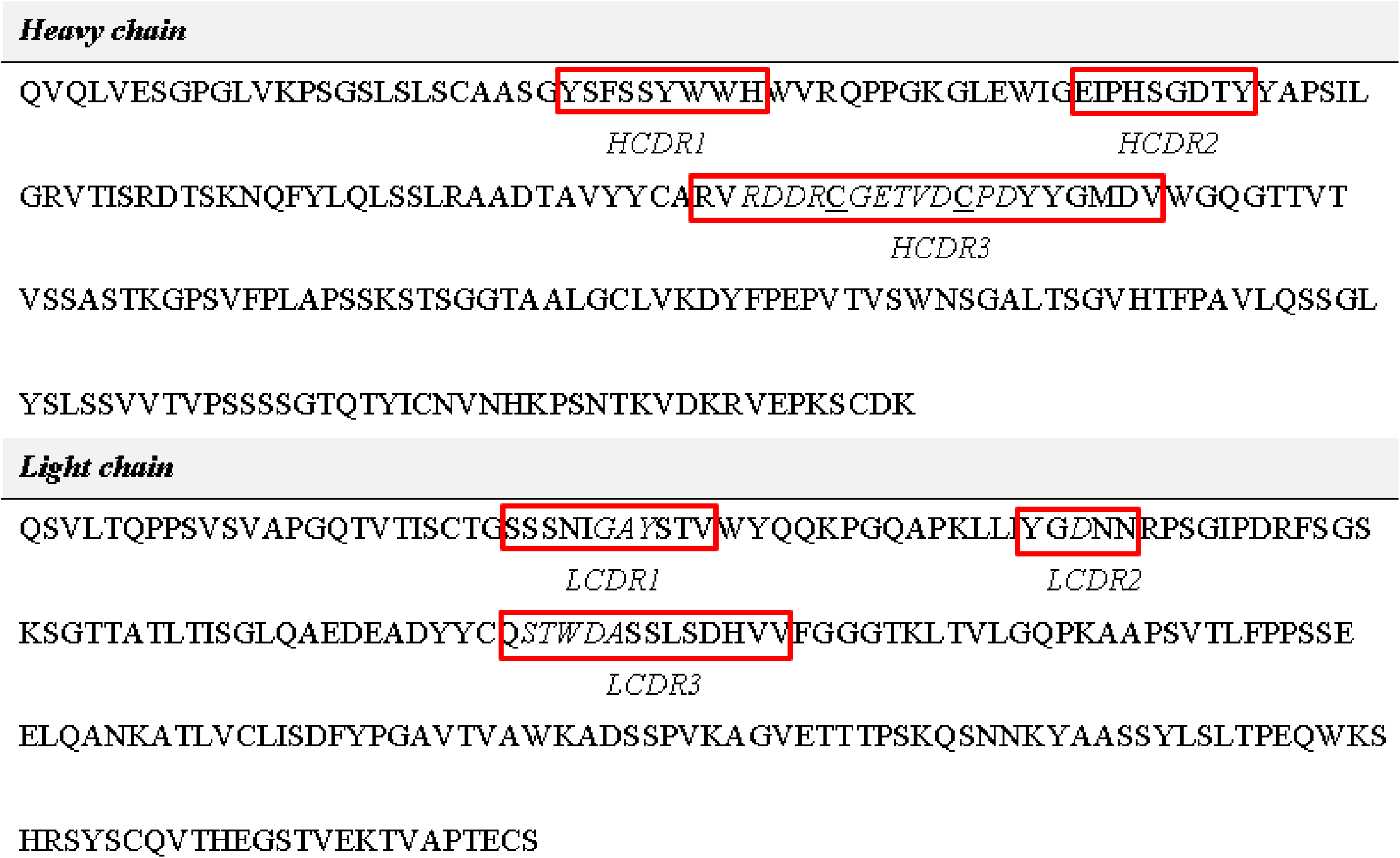
Proposed sequences for broad-spectrum Fab model against the H1 subtype of influenza A virus. *Italic amino acids* indicate the interaction with HA from templates, while <u>underlined amino acids</u> indicate the importance of the conformation of the HCDR3 loop’s conformation.

### 3D structure modeling of the Fab model gives an average −0.03 DOPE score, while CDR3 regions generated high energy

The sequences were then modeled and evaluated for the 3D structure using the MODELLER tool based on known multiple templates and loop refinement processes. As previous results showed the agreement between the 3D structure in different regions, the CDR loops are fundamental in this process, making direct interaction with HA proteins. A hundred structural models were generated for the Fab model; the top ten structural models’ energy profile was generated and plotted to analyze the stability of generated 3D structures (Fig 2). The residual DOPE scores of the top ten models of the proposed Fab were around −0.03, most of them gave lower DOPE score as low as −0.045. The structural model 28, which had a more stable energy profile than other models, gave the lowest global DOPE score further evaluated. However, residues at H/LCDR3 loops, the important interacted regions with RBS of HA protein, possess quite a high energy profile that would suggest bad modeled loops for a later experiment.[95] MODELLER’s comparative modeling does not reflect the dynamic of protein structure; thus, these DOPE scores only present the comparable property with other models.[83] Those high DOPE regions were then going under *ab-initio* loop refinement for another hundred outputs in each step. Each peak was done separately until the global DOPE score stable. Six steps of loop refinement were done for six peaks having a score equal to or higher than - 0.02. As a result, from step 4, the DOPE scores are not different within later steps. Thus, the structure at loop refinement step 03 was chosen for the subsequent analysis.

**Fig 2.**
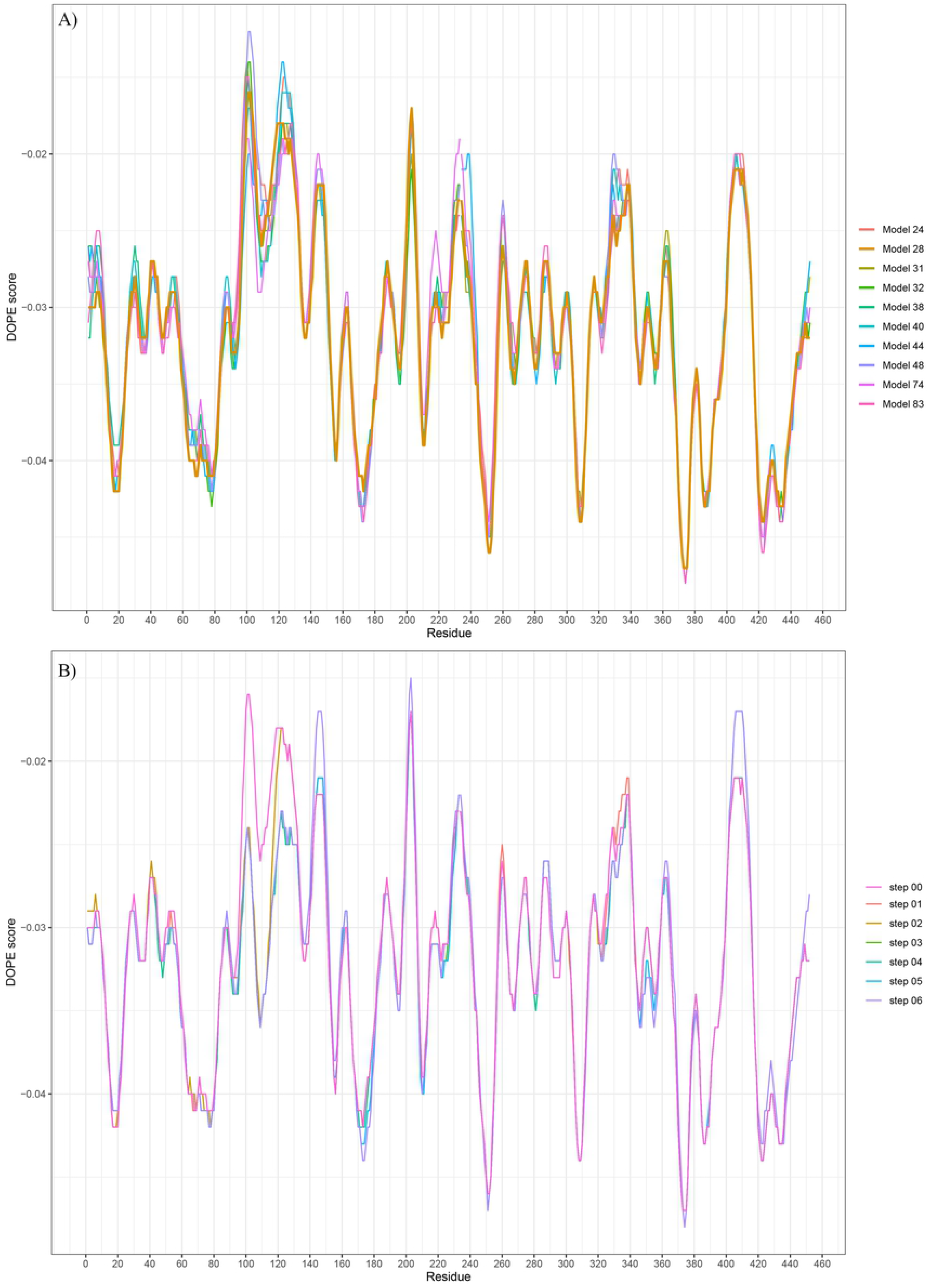
**Energy profile of the Fab model though different steps of 3D structure modeling** with plots of A) top ten models generated by the multiple-template process, and B) six steps of loop-refinement by the ab-initio method. The modeling and energy profiles were calculated using the MODELLER v9.21 tool, and the plots were generated using RStudio software.

### Broad-spectrum Fab models give good binding affinity on a wide range of H1N1 strains

Due to the 3D structure of the model constructed by a computational program, we used flexible docking between CDRs of Fab and RBS of HA for maximum results. As the protein in the biological environment can be flexible, especially the CDR loops, which have a high energy profile, could be fluctuated depending on the interaction, rigid docking is not suitable for this binding examination. Flexible docking of Fab templates and respective HA give results which is consistent with an original pose from experimental methods (Fig 3A). All the best cluster in each case has the lowest binding energy comparing with other clusters. Most of them have a binding position matching an original pose from X-ray crystallography, except complex systems 641 I-9 + SI-3/06 and H2227 + SI-3/06. While the original pose of H2227 with SI-3/06 can be found in cluster 1, 641 I-9 + SI-3/06 system does not show any sophisticated matching with original pose in flexible docking. In experimental results, 641 I-9 and SI-03/06 made contacts only through the heavy chain of Fab,[35] while in our docking experiment, the residues in both heavy and light chains were examined. Also, the 3D structure fetched from RCSB had missed some residues (only from residue 55 to 260 of HA head), making the docking incorrectly. In the interaction between H2227 and same influenza strain, when investing through X-ray, although the HCDR3 had contact with RBS of HA, this effect is little comparing with interactions forming between HA between HCDR1 and HCDR2.[32] Following the number of clusters generated by HADDOCK, 5J8FAB, and Nag-09N083/06 system has the most stable binding with the lowest number of clusters, and H1244 + Bei-262/95 system has varied binding poses with the highest number of clusters. Compared with the mean of the binding energy of all original-similar poses, the 2D1, CH65, and 6639 Fabs give good binding affinity with corresponding HA over other Fabs.

**Fig 3.**
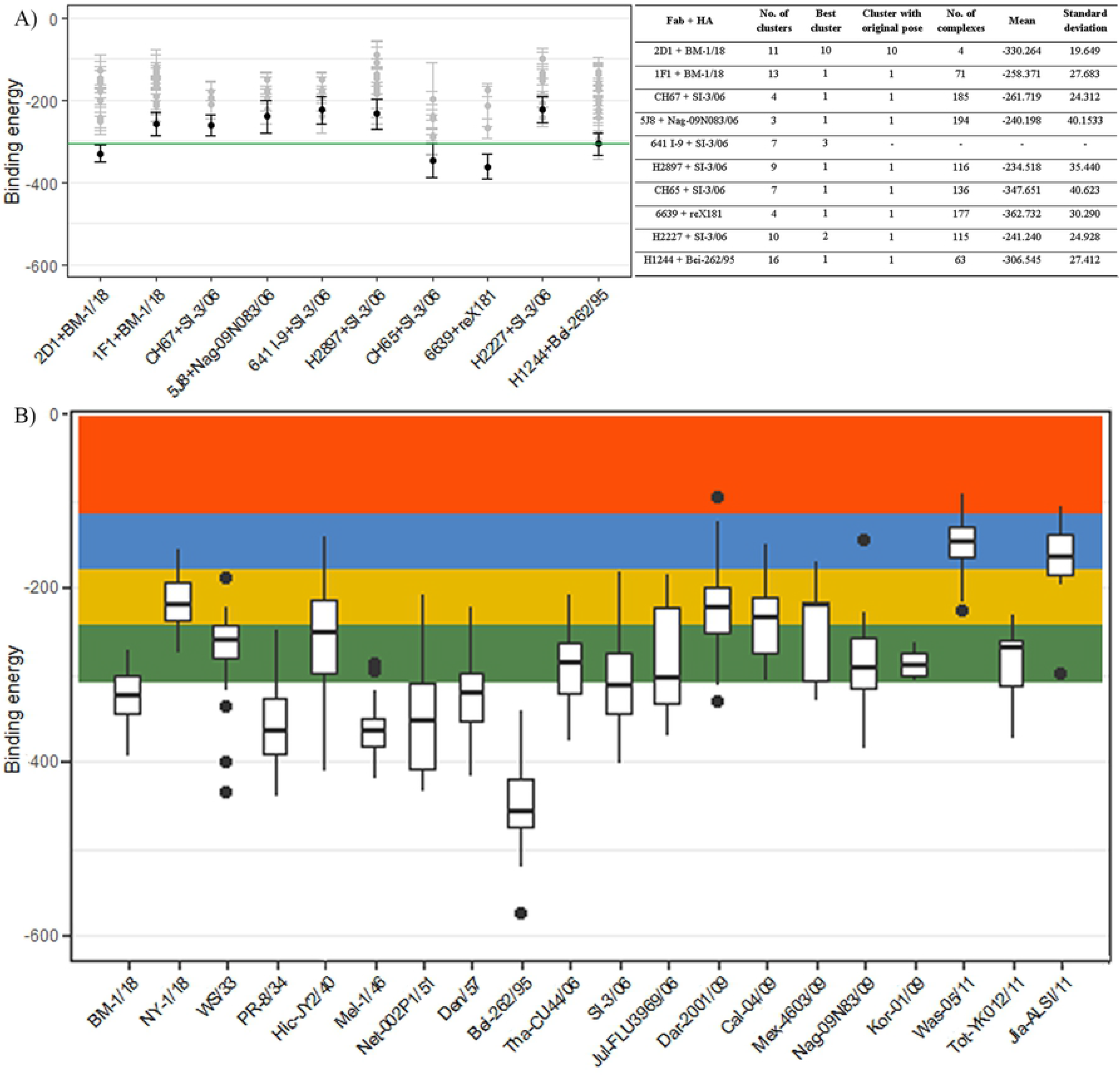
Binding energy of Fab RBS of HA protein of different strains of H1N1 influenza A virus. (A) The binding energy of different Fab templates on their corresponding HA was calculated by the HADDOCK tool. The best cluster of each case was expressed as black, while other clusters showed in gray, mean value of all complexes in original clusters was colored in green. Data were shown in the mean and standard deviation bars. (B) The binding energy of the proposed Fab model at RBS of HA protein of different H1N1 influenza A virus strains were shown as boxplot comparing with templates. Shaded error of binding energy of templated was indicated by colors: mean and green = 1s.d., yellow = 2s.d., blue = 3s.d., and red > 3s.d. The clustering of results was done by the HADDOCK tool based on interface Root-mean-square deviation (RMSD) and Fraction of common contacts (FCC).

The mean and standard deviation of binding energy from these original-matching poses was calculated and mapped on Fig 3B to evaluate the Fab model’s binding affinity with different strains of influenza A virus. There are 20 strains from 1918 to 2011 had been selected, not including the other seven mutated strains, and run with the Fab model in flexible docking. Similar to template analysis, the best cluster in each case was chosen for further analysis. The mean and standard deviation of binding energy was calculated and plotted in the same Fig 3. As a result, these complexes’ binding energy is agreed with a normal distribution (μ −308.563, δ 64.642) followed Fab templates when mean and standard deviation lay within 3 standard deviations of the mean. This result can be confirmed the broad-spectrum activity and binding effect of the model Fab on all 20 different H1N1 strains of influenza A virus. Among those 20 strains, the binding affinity of the Fab model and BM-1/18, PR-8/34, Mel-1/46, Net-002P1/51, Den/57, SI-3/06, and Jul-FLU3969/06 is higher than templated complexes. The research on these strains was conducted through pandemic 1918, and there is a long time for people developing immune system against these strains, the information gathered from these strain-against Fabs provides proper designation. While all strains from pandemic 2009 show a low binding affinity between the Fab model and HA.

The Fab model has the best binding affinity or lowest binding energy with Bei-262/95 strain, while it possesses high binding energy with Was-05/11 and Jia-ALSI/11, which are both from year-2011 strains. Although the Was-05/11 and Tot-YK012/11 have the same amino acid sequence at the head, their 3D structures at RBS have little difference (0.555 RMSD), which causes the difference in docking result between the Fab model and these strains. Meanwhile, Jia-ALSI/11 is an entirely new strain which is not related to other 2011 strains. The MSA of amino acid sequence at the head of these strains showed a distinguished Jiangsu strain in 2011, which explained the low binding affinity of Fab with this strain. There is no report on this strain, the work related to this published 3D structure on RCSB is still not published yet; thus, the Fab model did not have any residues information interacting with this strain. When observing the Minimum evolution phylogenetic tree constructed by MEGA-X, the Jia-ALSI/11 has the same origin as the 1918-pandemic strain, especially BM-1/18. It means this strain’s genetic material is still wandered around and had mutations to invade the immune system. A complement to this conclusion is the low binding affinity of the Fab model to this Jia-ALSI/11 strain. Meanwhile, the high binding energy between the Fab model and NY-1/18 can be explained by high mutated of this strain compared to BM-1/18.

In our flexible docking experiment, mutated BM-1/18, SI-3/06, and Cal-04/09 had been collected and run together with the wild-type strain (Fig 4). When comparing the HADDOCK score of the best cluster between wild-type and mutated strains, there is no significant difference between groups in two previous strains, while it is significantly different between wild-type and mutated types in Cal-04/09 strains. The HADDOCK score is recorded poorly in wild-type; thus, it is concluded that the interaction made by the Fab model and wild-type RBS is fewer than in mutated RBS. This result is consistent with the analysis of binding energy between them and confirms our conclusion. The Fab model had better affinity on mutated strains than wild type strain, which is essential in drug development against these viruses. Meanwhile, the binding affinity between the Fab model and other mutated BM-1/18 and SI-3/06 is different from wild type strain. The RBS in wild type made better interaction with the Fab model than the RBS in mutated strains. In BM-1/18, Lys’s adding at 130 in 130-loop of RBS made the loop longer; thus, it alternates some interaction, although it makes other types of interaction. This add130K deleted two salt bridges made between the Fab model with Lys219 and Asp222. In the SI-3/06 strain, the mutations are identified with three changes; only two are involved in the RBS region. However, when examining protein interaction, two T194K and R223Q mutations are not involved in the interaction between the Fab model and HA. The docking position could explain the significant difference in binding energy between wild type and mutation. As the scatter plot of binding energy and HADDOCK scores of complexes in the best cluster are shown, the Fab model’s pose on RBS of HA might be varied, and between wild type and mutation, there will be complexes with same binding poses.

**Fig 4.**
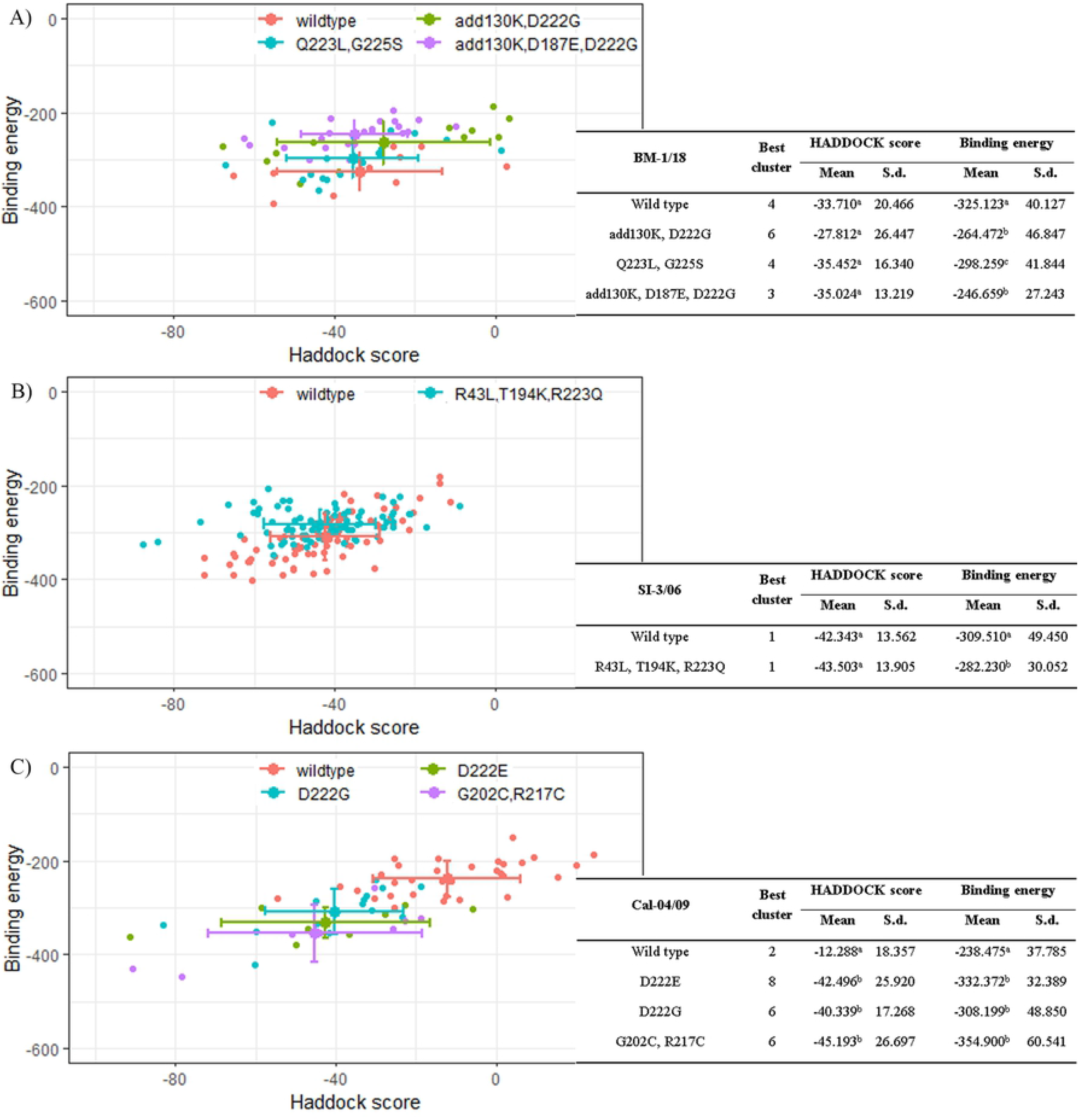
Binding energy and HADDOCK score between the proposed Fab model and the HA of influenza A virus. Wild type and different mutation types in strain A) BM-1/18, B) SI-3/06, and C) Cal-04/09 gave different effects in binding with the proposed Fab model. The plots were generated using RStudio software. RStudio calculated p-value by one-way ANOVA – between mean of each parameter of mutations within a strain, different alphabets stand for statistically significant difference (α=0.05).

### Lysine in 130-, 150-, and 220- loops play a vital role in forming electrostatic interaction between the Fab model and HA. Primary hydrophobic interaction formed between Fab model and HA made by Alanine, Leucine, and Valine in 130- loop and 190- helix

The best binding complex between the Fab model and each HA was investigated in protein-protein interaction using BIOVIA Discovery studio and Protein Interaction Calculator. This analysis will give the preliminary results to explain the difference in binding affinity between strains and mutations. All types of interaction were collected and put on the map to gather the information. The interaction between the Fab model and RBS of HA is mainly formed by hydrophobic interaction, hydrogen bond, electronic interaction, and pi-sulfur bond. There is only one pi-sulfur bond formed between Cys103H of Fab and Try150 of RBS on HA. Most strains have this interaction, except influenza A virus strains in the year 2011. However, the binding affinity with these strains is not hight; thus, this bond might not be a significant interaction. Hydrophobic interaction is mainly formed between the Fab model with 130- loop and 190- helix turn of RBS region. Especially, Lys130, Val132, Ala134, and Leu191 are found in most strains and made interaction with the Fab model. Ala141 of 130- loop also generate hydrophobic interaction with the Fab model. This amino acid is found in 1918-, 2009- and 2011 strains, while in another influenza A strains, this Ala141 is alternated by Lysine or Glutamic acid, which does not form hydrophobic bonding. Similarly, Ala186 in 2009- and 2011 strains are recorded for forming this type of interaction with the Fab model. Talking about hydrogen bonds, which is considered as the main interaction between protein and protein, both conventional and carbon-hydrogen bonds were observed through all loops and helix forming RBS of HA. The interaction between amino acids of the Fab model is throughout the length of RBS, mainly formed by Threonine, Valine, Lysine, Alanine, Glycine, Serine, Aspartic acid, and Glutamine. The number of hydrogen bonds is not significantly different between wild type and mutated strains; the mutated amino acids could also form the hydrogen bonding between the Fab model and HA. Observing the salt bridge made between the Fab model and RBS of HA, it is seen that most of the interaction is formed at 220- loop. While in 1918 strains, Glu127 makes a salt bridge with Arg99^H^, Glu105^H^, and Lys67^L^, in other strains, salt bridges are made by Lys130 with Asp101^H^, Asp108^H^, or Lys67^L^. Other salt bridges can be listed as formed Lysine at 130-, 150- and 220- loop of HA. At 220- loop, salt bridges are also found by forming Asp222 and Glu224 of 1918, 2006, and 2009, 2011 strains, respectively. Notably, only strains from 1934 to 2006 have amino acids at 190- helix forming salt bridges with the Fab model. Electronic interactions, including attractive charge and pi-cation interaction, are also found between the Fab model and RBS of HA. Lysine and Glutamine form most of them throughout the sequence.

## Conclusions

This study lays the groundwork for developing a universal therapeutic vaccine, especially for the influenza A virus, by bioinformatic approach. The proposed Fab model of broad-spectrum antibody targeting HA of different H1N1 Influenza A strains had preliminarily shown the broad-reactive property against RBS of HA. It shows a positive view in applying bioinformatics in dealing with emergent pandemics. The interaction between the Fab model and HA can be sketchy when protein-protein interaction is analyzed by software. The correct binding position should be investigated deeply with simulation, and the critical interaction could be obtained by steered molecular dynamics to explain the different binding affinity. This information will be useful to refine the Fab model, which will eliminate or replace the weak interaction with other amino acids. Furthermore, the Fab model should be examined the physical properties to make sure that it is suitable for the therapeutic vaccine and can pass phase 1 of a clinical test. In the next paper, these results will be reported, and further information will be provided for a better rational therapeutic vaccine design.

## Acknowledgment

We thank Mr. Cong T. Nguyen for his help in data analysis and presentation by RStudio software for this manuscript. This work was supported by the NAFOSTED (The National Foundation for Science and Technology) under Grant No. 108.06-2017.332 and the Domestic Master/ PhD Scholarship Programme of Vingroup Innovation Foundation.

## Author contributions

P-C.D. and L.L. designed research; P-C.D., T.H.N. and U.T.M.V. collected data and performed research; P-C.D. analyzed data; P-C.D. and L.L. wrote the paper.

The authors declare no conflict of interest.

## REFERENCES

1. Is the world ready to respond to the next influenza pandemic? Exploring Lessons Learned from a Century of Outbreaks: Readiness for 2030; 2019: The National Academies Press.

2. Taubenberger, JK, Morens, DM. 1918 Influenza: the Mother of all pandemics. Emerging Infectious Diseases. 2006;12(1):8. doi: 10.3201/eid1201.050979.

3. Stuart-Harris, CH, Schild, GC. Influenza: the viruses and the disease.: Edward Arnold; 1958.

4. Cox, NJ, Subbarao, K. Global epidemiology of influenza: Past and Present. Annual Review of Medicine. 2000;51:407–21.

5. Jong, JCFD, Claas, ECJ, Osterhaus, ADME, Webster, RG, Lim, WL. A pandermic warning? Nature. 1997;389(554):1.

6. Lee, N, Hui, D, Wu, A, Chan, P, Cameron, P, Joynt, GM, et al. A major ourbreak of Severe acute raspiratoty syndrome in Hong Kong. The New England Journal of Medicine. 2003;348(20):9.

7. Monto, AS, Webster, RG. Influenza pandemics: History and lessons learned. In: Webster, RG, Monto, AS, Braciale, TJ, Lamb, RA, editors. Textbook of Influenza. 2nd ed: Wiley Blackwell; 2013. p. 14.

8. Prevention and control of seasonal influenza with vaccines: Recommendations of the Advisory Committee on Immunization Practices (ACIP)— United, States, 2019-20 Centers for Disease Control and, Prevention, National Center for Immunization and Respiratory Diseases (NCIRD)2019 [16 Feb 2020]. Available from: https://www.cdc.gov/flu/professionals/acip/summary/summary-recommendations.htm#anchor-star.

9. Jennings, GT, Bachmann, MF. Immunotherapies: cause for measured optimism. Drug Discovery Today. 2002;7:994–6.

10. Bangaru, S, Lang, S, Schotsaert, M, Vanderven, HA, Zhu, X, Kose, N, et al. A site of vulnerability on the influenza virus hemagglutinin head domain trimer interface. Cell. 2019;177:1136–52.

11. Bangaru, S, Zhang, H, Gilchuk, IM, Voss, TG, Irving, RP, Gilchuk, P, et al. A multifunctional human monoclonal neutralizing antibody that targets a unique conserved epitope on influenza HA. Nature Communications. 2018;9:15.

12. Hong, M, Lee, PS, Hoffman, RMB, Zhu, X, Krause, JC, Laursen, NS, et al. Antibody recognition of the pandemic H1N1 influenza virus hemagglutinin receptor binding site. Journal of Virology. 2013;87:12471–80.

13. Sevy, AM, Wu, NC, Gilchuk, IM, Parrish, EH, Burger, S, Yousif, D, et al. Multistate design of influenza antibodies improves affinity and breadth against seasonal viruses. Proceedings of the National Academy of Sciences of the United States of America. 2018:6.

14. Tsibane, T, Ekiert, DC, Krause, JC, Martinez, O, Jr. JEC, Wilson, IA, et al. Influenza human monoclonal antibody 1F1 interacts with thre major antigenic sites and residues mediating human receptor specificity in H1N1 viruses. PLOS Pathogens. 2012;8(12):e1003067.

15. Turner, HL, Pallesen, J, Lang, S, Bangaru, S, Urata, S, Li, S, et al. Potent anti-influenza H7 human monoclonal antibody induces separation of hemagglutinin receptor-binding head domains. PLOS Biology. 2019:23. doi: 10.1371/journal.pbio.3000139.

16. Winarski, KL, Thornburg, NJ, Yu, Y, Sapparapu, G, Jr. JEC, Spiller BW. Vaccine-elicited antibody that neutralizes H5N1 influenza and variants binds the receptor site and polymorphic sites. Proceedings of the National Academy of Sciences of the United States of America. 2015;112:9346–51.

17. Xu, R, Ekiert, DC, Krause, JC, Hai, R, Jr. JEC, Wilson IA. Structural basis of preexisting immunity to the 2009 H1N1 pandemic influenza virus. Science. 2010;328:357–60.

18. Xu, R, Krause, JC, McBride, R, Paulson, JC, James, E. Crowe, J, Wilson, IA. A recurring motif for antibody recognition of the receptorbinding site of influenza hemagglutinin. Nature Structural & Molecular Biology. 2013;20:363–71.

19. Dreyfus, C, Ekiert, DC, Wilson, IA. Structure of a classical broadly neutralizing stem antibody in complex with a pandemic H2 influenza virus hemagglutinin. Journal of Virology. 2013;87:7149–54.

20. Dreyfus, C, Laursen, NS, Kwaks, T, Zuijdgeest, D, Khayat, R, Ekiert, DC, et al. Highly conserved protective epitopes on influenza B viruses. Science. 2012;337:1343–8.

21. Ekiert, DC, Bhabha, G, Elsliger M-A, Friesen, RHE, Jongeneelen, M, Throsby, M, et al. Antibody recognition of a highly conserved influenza virus epitope. Science. 2009;324:246–51.

22. Ekiert, DC, Friesen, RHE, Bhabha, G, Kwaks, T, Jongeneelen, M, Yu, W, et al. A highly conserved neutralizing epitope on group 2 influenza A viruses. Science. 2011;333.

23. Friesen, RHE, Lee, PS, Stoop, EJM, Hoffman, RMB, Ekiert, DC, Bhabha, G, et al. A common solution to group 2 influenza virus neutralization. Proceedings of the National Academy of Sciences of the United States of America. 2014;111:445–50.

24. Lang, S, Xie, J, Zhu, X, Wu, NC, Lerner, RA, Wilson, IA. Antibody 27F3 broadly targets influenza A group 1 and group 2 hemagglutinins through a further variation in V_H_1-69 antibody orientation on the HA stem. Cell Reports. 2017;20:2935–43.

25. Lee, PS, Ohshima, N, Stanfield, RL, Yu, W, Iba, Y, Okuno, Y, et al. Receptor mimicry by antibody F045-092 facilitates universal binding to the H3 subtype of influenza virus. Nature Communications. 2014;5:9.

26. Lee, PS, Yoshida, R, Ekiert, DC, Sakai, N, Suzuki, Y, Takada, A, et al. Heterosubtypic antibody recognition of the influenza virus hemagglutinin receptor binding site enhanced by avidity. Proceedings of the National Academy of Sciences of the United States of America. 2012;109:17040–5.

27. Wu, NC, Grande, G, Turner, HL, Ward, AB, Xie, J, Lerner, RA, et al. In vitro evolution of an influenza broadly neutralizing antibody is modulated by hemagglutinin receptor specificity. Nature Communications. 2017;8.

28. Wu, NC, Yamayoshi, S, Ito, M, Uraki, R, Kawaoka, Y, Wilson, IA. Recurring and adaptable binding motifs in broadly neutralizing antibodies to influenza virus are encoded on the D3-9 segment of the Ig gene. Cell Host & Microbe. 2018:569–78.

29. Wyrzucki, A, Dreyfus, C, Kohler, I, Steck, M, Wilson, IA, Hangartner, L. Alternative recognition of the conserved stem epitope in influenza A virus hemagglutinin by a V_H_3-30-encoded heterosubtypic antibody. Journal of Virology. 2014;88:7083–92.

30. Zhu, X, Guo Y-H, Jiang, T, Wang Y-D, Chan K-H, Li X-F, et al. A unique and conserved neutralization epitope in H5N1 influenzaviruses identified by an antibody against the A/Goose/Guangdong/1/96 hemagglutinin. Journal of Virology. 2013;87:12619–35.

31. Liu, Y, Pan, J, Jenni, S, Raymond, DD, Caradonna, T, Do, KT, et al. CryoEM structure of an influenza virus receptor-binding site antibody-antigen interface. Journal of Molecular Biology. 2017;429:1829–39.

32. McCarthy, KR, Raymond, DD, Do, KT, Schmidt, AG, Harrison, SC. Affinity maturation in a human humoral response to influenza hemagglutinin. Proceedings of the National Academy of Sciences of the United States of America. 2019;116:26745–51. doi: 10.1073/pnas.1915620116.

33. McCarthy, KR, Watanabe, A, Kuraoka, M, Do, KT, McGee, CE, Sempowski, GD, et al. Memory B cells that cross-react with group 1 and group 2 influenza a viruses are abundant in adult human repertoires. Immunity. 2018;48:174–84.

34. Raymond, DD, Bajic, G, Ferdman, J, Suphaphiphat, P, Settembre, EC, Moody, MA, et al. Conserved epitope on influenza-virus hemagglutinin head defined by a vaccine-induced antibody. Proceedings of the National Academy of Sciences of the United States of America. 2017;115:168–73.

35. Schmidt, AG, Therkelsen, MD, Stewart, S, Kepler, TB, Liao H-X, Moody, MA, et al. Viral receptor-binding site antibodies with diverse germline origins. Cell. 2015;161:1–9.

36. Schmidt, AG, Xu, H, Khan, AR, O’Donnell, T, Khurana, S, King, LR, et al. Preconfiguration of the antigen-binding site during affinity maturation of a broadly neutralizing influenza virus antibody. Proceedings of the National Academy of Sciences of the United States of America. 2013;110(1):264–9.

37. Watanabe, A, McCarthy, KR, Kuraoka, M, Schmidt, AG, Adachi, Y, Onodera, T, et al. Antibodies to a conserved influenza head interface epitope protect by an IgG subtype-dependent mechanism. Cell. 2019;177:1124–35.

38. Whittle, JRR, Zhang, R, Khurana, S, King, LR, Manischewitz, J, Golding, H, et al. Broadly neutralizing human antibody that recognizes the receptor-binding pocket of influenza virus hemagglutinin. Proceedings of the National Academy of Sciences of the United States of America. 2011;108:14216–21.

39. Bajic, G, Maron, MJ, Adachi, Y, Onodera, T, McCarthy, KR, McGee, CE, et al. Influenza antigen engineering focuses immune responses to a subdominant but broadly protective viral epitope. Cell Host & Microbe. 2019;25:827–35.

40. Benton, DJ, Nans, A, Calder, LJ, Turner, J, Neu, U, Lin, YP, et al. Influenza hemagglutinin membrane anchor. Proceedings of the National Academy of Sciences of the United States of America. 2018:6.

41. Corti, D, Voss, J, Gamblin, SJ, Codoni, G, Macagno, A, Jarrossay, D, et al. A neutralizing antibody selected from plasma cells that binds to group 1 and group 2 influenza A hemagglutinins. Science. 2011;333:850–6.

42. Kallewaard, NL, Corti, D, Collins, PJ, Neu, U, McAuliffe, JM, Benjamin, E, et al. Structure and function analysis of an antibody recognizing all influenza A subtypes. Cell. 2016;166:596–608.

43. Xiong, X, Corti, D, Liu, J, Pinna, D, Foglierini, M, Calder, LJ, et al. Structures of complexes formed by H5 influenza hemagglutinin with a potent broadly neutralizing human monoclonal antibody. Proceedings of the National Academy of Sciences of the United States of America. 2015;112:9430–5.

44. Wang, P, Zuo, Y, Sun, J, Zuo, T, Zhang, S, Guo, S, et al. Structural and functional definition of a vulnerable site on the hemagglutinin of highly pathogenic avian influenza A virus H5N1. Journal of Biological Chemistry. 2019:25.

45. Zuo, T, Sun, J, Wang, G, Jiang, L, Zuo, Y, Li, D, et al. Comprehensive analysis of antibody recognition in convalescent humans from highly pathogenic avian influenza H5N1 infection. Nature Communications. 2015;6:12.

46. Zuo, Y, Wang, P, Sun, J, Guo, S, Wang, G, Zuo, T, et al. Complementary recognition of the receptor-binding site of highly pathogenic H5N1 influenza viruses by two human neutralizing antibodies. Journal of Biological Chemistry. 2018;293:16503–17.

47. Joyce, MG, Wheatley, AK, Thomas, PV, Chuang G-Y, Soto, C, Bailer, RT, et al. Vaccine-induced antibodies that neutralize group 1 and 2 influenza a viruses. Cell. 2016;166:609–23.

48. Wang, W, Sun, X, Li, Y, Su, J, Ling, Z, Zhang, T, et al. Human antibody 3E1 targets the HA stem region of H1N1 and H5N6 influenza A viruses. Nature Communications. 2016;7.

49. Xiao, H, Guo, T, Yang, M, Qi, J, Huang, C, Hong, Y, et al. Light chain modulates heavy chain conformation to change protection profile of monoclonal antibodies against influenza A viruses. Cell Discovery. 2019;5:16.

50. Padlan, EA, Abergel, C, Tipper, JP. Identification of specificity-determining residues in antibodies. Research Communications. 1995:7. doi: 10.1096/fasebj.9.1.7821752.

51. M. MacCallum R, C.R. Martin A, M. Thornton J. Antibody-antigen interactions: Contact analysis and binding site topography. Journal of Molecular Biology. 1996;262:732–45. doi: 10.1006/jmbi.1996.0548.

52. Kunik, V, Ashkenazi, S, Ofran, Y. Paratome: an online tool for systemati identification of antigen-binding regions in antibodies based on sequence or structure. Nucleic Acids Research. 2012;40:W521–4.

53. Schroeder, HW, Cavacini, L. Structure and function of immunoglobulins. Journal of Allergy and Clinical Immunology. 2010;125(2):S41–S52. doi: 10.1016/j.jaci.2009.09.046.

54. Smith, KA, Nelson, PN, Warren, P, Astley, SJ, Murray, PG, Greenman, J. Demystified… recombinant antibodies. Journal of Clinical Pathology. 2004;57:912–7. doi: 10.1136/jcp.2003.014407.

55. Raymond, DD, Stewart, SM, Lee, J, Ferdman, J, Bajic, G, Do, KT, et al. Influenza immunization elicits antibodies specific for an egg-adapted vaccine strain. Nature Medicine. 2016;22:1465–9.

56. Ni, F, Kondrashkina, E, Wang, Q. Determinant of receptor-preference switch in influenza hemagglutinin. Virology. 2018;513:98–107. doi: 10.1016/j.virol.2017.10.010.

57. Zhang, W, Shi, Y, Qi, J, Gao, F, Li, Q, Fan, Z, et al. Molecular basis of the receptor binding specificity switch of the hemagglutinins from both the 1918 and 2009 pandemic influenza A viruses by a D225G substitution. Journal of Virology. 2013;87(10):5949–58. doi: 10.1128/JVI.00545-13.

58. Tsibane, T, Ekiert, DC, Krause, JC, Martinez, O, Jr, JEC, Wilson, IA, et al. Influenza human monoclonal antibody 1F1 interacts with three major antigenic sites and residues mediating human receptor specificity in H1N1 viruses. PLOS Pathogens. 2012;8(12):e1003067. doi: 10.1371/journal.ppat.1003067.

59. Conrady, DG, Dranow, DM, Calhoun, B, Lorimer, DD, Horanyi, PS, Edwards, TE. Crystal structure of InvbP.18715.a.KN11: Influenza hemagglutinin from strain A/Almaty/32/1998. To be published.

60. Chevalier, A, Silva D-A, Rocklin, GJ, Hicks, DR, Vergara, R, Murapa, P, et al. Massively parallel de novo protein design for targeted therapeutics. Nature. 2017;550(7674):74–9. doi: 10.1038/nature23912.

61. Conrady, DG, Mayclin, SJ, Calhoun, B, Lorimer, DD, Horanyi, PS, Edwards, TE. Crystal structure of Influenza hemagglutinin from strain A/Hickox/JY2/1940. To be published.

62. Conrady, DG, Yano, JK, Calhoun, B, Lorimer, DD, Horanyi, PS, Edwards, TE. Crystal structure of Influenza hemagglutinin from strain A/Melbourne/1/1946(H1N1). To be published.

63. Conrady, DG, Calhoun, B, Lorimer, DD, Horanyi, PS, Edwards, TE. Crystal structure of InvbM.18715.a.KN11: Influenza hemagglutinin from strain A/Netherlands/002P1/1951. To be published.

64. Edwards, TE, Conrady, DG, Horanyi, PS, Lorimer, DD, Disease, SSGCfI. Crystal structure of hemagglutinin from H1N1 Influenza A virus A/Denver/57 bound to the C05 antibody. To be published.

65. McCarthy, KR, Raymond, DD, Do, KT, Schmidt, AG, Harrison, SC. Affinity maturation in a human humoral response to influenza hemagglutinin. Proceedings of the National Academy of Sciences of the United States of America. 2019;116(52):26744–51. doi: 10.1073/pnas.1915620116.

66. Cho, KJ, Lee J-H, Hong, KW, Kim S-H, Park, Y, Lee, JY, et al. Insight into structural diversity of influenza virus haemagglutinin. Journal of General Virology. 2013;94(Pt 8):1712–22. doi: 10.1099/vir.0.051136-0.

67. Raymond, DD, Bajic, G, Ferdman, J, Suphaphiphat, P, Settembre, EC, Moody, MA, et al. Conserved epitope on influenza-virus hemagglutinin head defined by a vaccine-induced antibody. Proceedings of the National Academy of Sciences of the United States of America. 2018;115(1):168–73. doi: 10.1073/pnas.1715471115.

68. Dongen, MJPv, Kadam, RU, Juraszek, J, Lawson, E, Brandenburg, B, Schmitz, F, et al. A small-molecule fusion inhibitor of influenza virus is orally active in mice. Science. 2019;363(6431):eaar6221. doi: 10.1126/science.aar6221.

69. Dong, J, Sevy, AM, Crowe, JE. High resolution crystal structure of the hemagglutinin H1 head domain of influenza A virus Solomon Islands. To be published.

70. Yang, H, Carney, P, Stevens, J. Structure and receptor binding properties of a pandemic H1N1 virus hemagglutinin. PLoS Currents. 2010:RRN1152. doi: 10.1371/currents.RRN1152.

71. Xu, R, Ekiert, DC, Krause, JC, Hai, R, Jr, JEC, Wilson, IA. Structural basis of preexisting immunity to the 2009 H1N1 pandemic influenza virus. Science. 2010;328(5976):357–60. doi: 10.1126/science.1186430.

72. Xuan, C, Shi, Y, Qi, J, Zhang, W, Xiao, H, Gao, GF. Structural vaccinology: structure-based design of influenza A virus hemagglutinin subtype-specific subunit vaccines. Protein & Cell. 2011;2(12):997–1005. doi: 10.1007/s13238-011-1134-y.

73. Xu, R, McBride, R, Nycholat, CM, Paulson, JC, Wilson, IA. Structural characterization of the hemagglutinin receptor specificity from the 2009 H1N1 influenza pandemic. Journal of Virology. 2012;86(2):982–90. doi: 10.1128/JVI.06322-11.

74. DuBois, RM, Aguilar-Yañez, JM, Mendoza-Ochoa, GI, Oropeza-Almazán, Y, Schultz-Cherry, S, Alvarez, MM, et al. The receptor-binding domain of influenza virus hemagglutinin produced in Escherichia coli folds into its, native, immunogenic, structure. Journal of Virology. 2011;85(2):865–72. doi: 10.1128/JVI.01412-10.

75. Hong, M, Lee, PS, Hoffman, RMB, Zhu, X, Krause, JC, Laursen, NS, et al. Antibody recognition of the pandemic H1N1 influenza virus hemagglutinin receptor binding site. Journal of Virology. 2013;87(22):12471–80. doi: 10.1128/JVI.01388-13.

76. Yang, H, Chang, JC, Guo, Z, Carney, PJ, Shore, DA, Donis, RO, et al. Structural stability of influenza A(H1N1)pdm09 virus hemagglutinins. Journal of Virology. 2014;88(9):4828–38. doi: 10.1128/JVI.02278-13.

77. Conrady, DG, Fox III, D, Horanyi, PS, Lorimer, DD, Edwards, TE. Crystal structure of InvbI.18715.a.KN11: Influenza hemagglutinin from strain A/Jiangsu/ALSI/2011. To be published.

78. Kumar, S, Stecher, G, Li, M, Knyaz, C, Tamura, K. MEGA X: Molecular Evolutionary Genetics Analysis across computing platforms. Molecular Biology and Evolution. 2018;35(6):1547–9. doi: 10.1093/molbev/msy096.

79. Kaas, Q, Ruiz, M, Lefranc M-P. IMGT/3Dstructure-DB and IMGT/StructuralQuery, a database and a tool for, immunoglobulin, T cell receptor and MHC structural data. Nucleic Acids Research. 2004;32:D208–10.

80. Ehrenmann, Fo, Kaas, Q, Lefranc M-P. IMGT/3Dstructure-DB and IMGT/DomainGapAlign: a database and a tool for immunoglobulins or, antibodies, T cell, receptors, MHC, IgSF and MhcSF. Nucleic Acids Research. 2010;38:D301–7.

81. Ehrenmann, F, Lefranc M-P. IMGT/3Dstructure-DB: Querying the IMGT database for 3D structures in immunology and immunoinformatics (Ig or, antibodies, TR, MH, RPI, and FPIA). Cold Spring Harbor Protocols. 2011:750–61.

82. Pettersen, EF, Goddard, TD, Huang, CC, Couch, GS, Greenblatt, DM, Meng, EC, et al. UCSF Chimera—A visualization system for exploratory research and analysis. Journal of Computational Chemistry. 2004;25:1605–12. doi: 10.1002/jcc.20084.

83. Eswar, N, Eramian, D, Webb, B, Shen M-Y, Sali A. Protein structure modeling with MODELLER. In: Kobe, B, Guss, M, Huber, T, editors. Structural Proteomics-High-throughput Methods. Methods in Molecular Biology™. 426: Humana Press; 2008. p. 145–59.

84. Shen M-y, Sali A. Statistical potential for assessment and prediction of protein structures. Protein Science. 2006;15:2507–24. doi: 10.1110/ps.062416606.

85. Vries, SJd, Dijk, Mv, Bonvin, AMJJ. The HADDOCK web server for data-driven biomolecular docking. Nature Protocols. 2010;5:883–97. doi: 10.1038/nprot.2010.32.

86. Zundert, GCPv, Rodrigues, JPGLM, Trellet, M, Schmitz, C, Kastritis, PL, Karaca, E, et al. The HADDOCK2.2 web server: User-friendly integrative modeling of biomolecular complexes. Journal of Molecular Biology. 2016;428(4):720–5. doi: 10.1016/j.jmb.2015.09.014.

87. Tina, KG, Bhadra, R, Srinivasan, N. PIC: Protein Interactions Calculator. Nucleic Acids Research. 2007;35(Web server):W473–W6.

88. Sinz A. Chemical cross-linking and mass spectrometry to map three-dimensional protein structures and protein-protein interactions. Mass Spectrometry Reviews. 2006;25:663–82.

89. Levy, Y, Cho, SS, Onuchic, JN, Wolynes, PG. A survey of flexible protein binding mechanisms and their transition states using native topology based energy landscapes. Journal of Molecular Biology. 2005;346:1121–45.

90. Viguera A-R, Serrano L. Loop, length, intramolecular diffusion and protein folding. Nature Structural & Molecular Biology. 1997;4:939–46. doi: 10.1038/nsb1197-939.

91. Nagi, AD, Regan, L. An inverse correlation between loop length and stability in a four-helix-bundle protein. Folding and Design. 1997;2(1):67–75. doi: 10.1016/S1359-0278(97)00007-2.

92. Bankovich, AJ, Raunser, S, Juo, ZS, Walz, T, Davis, MM, Garcia, KC. Structural insight into pre-B cell receptor function. Science. 2007;316(5822):291–4. doi: 10.1126/science.1139412.

93. Le, TH, Nguyen, NTB. Evolutionary dynamics of highly pathogenic avian influenza A/H5N1 HA clades and vaccine implementation in Vietnam. Clinical and Experimental Vaccine Research. 2014;3:117–27.

94. Lee C-C, Chih-YaYang, Lin L-L, Ko T-P, Chang AH-L, Chang SS-C, et al. An efective neutralizing antibody against infuenza virus H1N1 from human B cells. Scientific Reports. 2019;9:11.

95. Webb, B, Sali, A. Comparative protein structure modeling using MODELLER. Current Protocols in Bioinformatics 2016;56:5.6.1–5.6.37. doi: 10.1002/cpbi.3.

